# Association between breeding-habitat bacterial communities and oviposition behavior of *Aedes aegypti:* Insights from arboviral outbreak-affected areas of Pune, Western India

**DOI:** 10.64898/2026.10.01.755851

**Authors:** Shalini Singh, Manas Raikar, Shruti Dongardive, Ashwini Ramdasi, Surendra Kumar, Sachin Dhaigude, Pratap Kadam, Rajlakshmi Viswanathan, Amit Yadav, Krishanpal Karmodiya, Kavita S. Lole, Kaushik Banerjee, Sreelakshmi PR, Tanwee Das De

## Abstract

*Aedes aegypti*, an important vector of several arboviral diseases, is highly adapted to human-associated environments and commonly exploits anthropogenic water-holding containers for breeding. Understanding the ecological factors governing oviposition-site selection is essential for developing sustainable vector control strategies. Previous studies have established that microbial communities in breeding waters influence mosquito oviposition-site selection through the production of volatile organic compounds. Importantly, rapid urbanization in India has increased the availability of artificial breeding habitats for *Ae. aegypti*, potentially facilitating the transmission and persistence of arboviral diseases. However, the composition of bacterial communities in these larval habitats and their potential role in shaping the oviposition behavior of *Ae. aegypti* remain poorly characterized in the Indian context. Here, we investigated the bacterial diversity and community composition of *Aedes* larval habitats during the 2024 Zika and chikungunya outbreaks in Pune, India. Water samples, collected from artificial urban containers and natural freshwater habitats, were subjected to 16S rRNA amplicon sequencing. Artificial breeding habitats exhibited significantly higher bacterial diversity and were enriched with *Enterobacter* spp. compared with natural habitats. Oviposition bioassays further demonstrated a strong preference of gravid *Ae. aegypti* females for *Enterobacter*-conditioned water. GC–MS analysis of bacterial suspension identified indole as a prominent volatile metabolite associated with *Enterobacter*, suggesting its possible role in mediating oviposition site selection. Together, these findings provide insights into the microbial and chemical ecology of *Ae. aegypti* breeding habitats and establish a basis for investigating microbial volatile cues as potential targets for vector surveillance and environmentally sustainable attract-and-kill strategies.

**Graphical Abstract:** 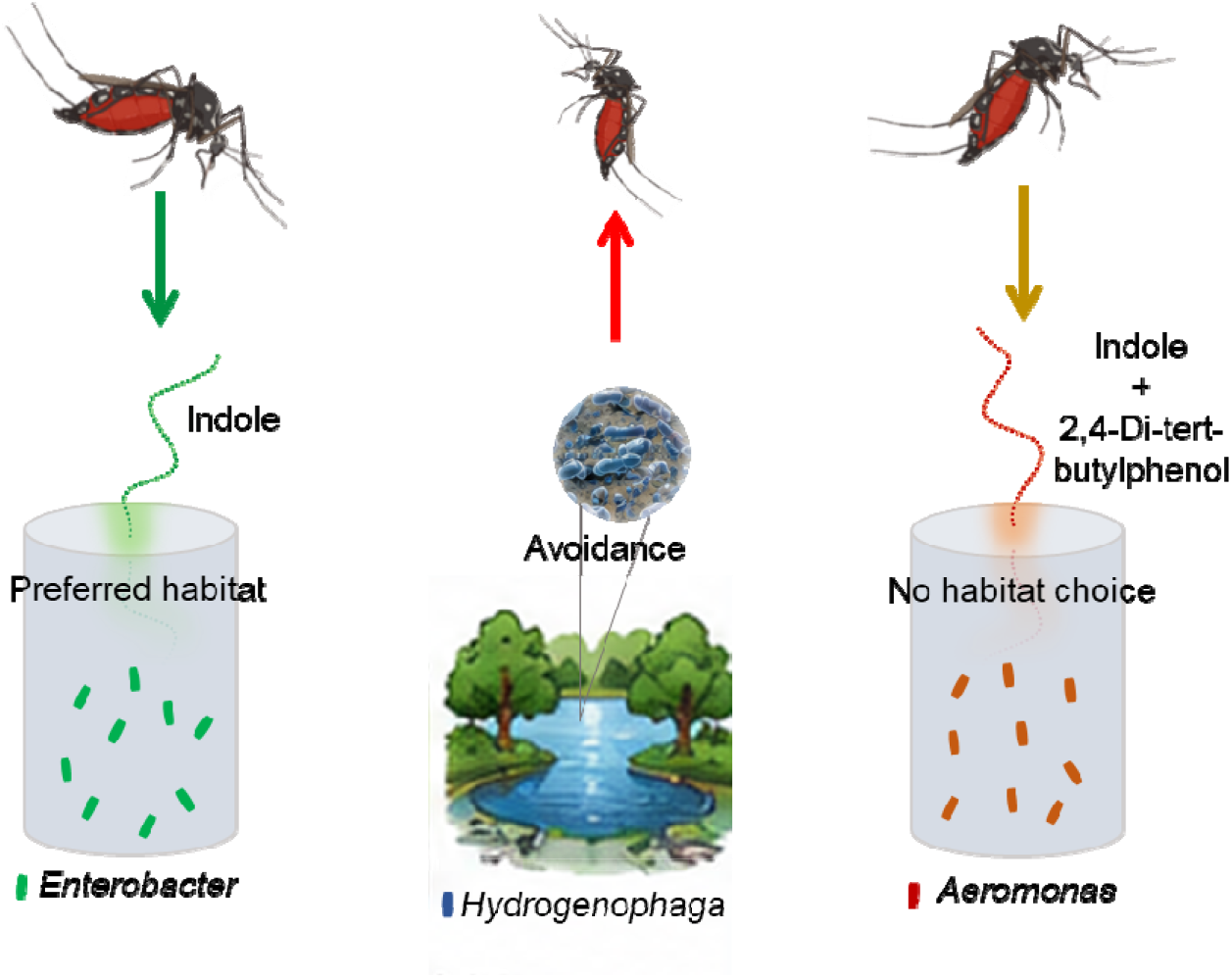

## Introduction

India remains one of the major contributors to the global burden of mosquito-borne diseases, including dengue, malaria, chikungunya, and filariasis (1). The vast geographical expanse, coupled with diverse ecological settings, climatic zones, and sociocultural heterogeneity, results in region-specific epidemiological patterns of vector-borne diseases (2). Rapid population growth and economic development have accelerated urbanization in recent decades. This has transformed land use, expanded construction and industrialization, and led to unplanned settlements, collectively fueling unprecedented arboviral transmission across India and South Asia (3).

Within this context, *Aedes aegypti* has demonstrated exceptional adaptability to urban environments, efficiently exploiting human hosts for blood meals and utilizing artificial water-holding containers as breeding sites (4). This adaptability has established the species as a persistent and significant public health threat. Thus, there is a pressing need for innovative and sustainable interventions against *Ae. aegypti*. Strategies focusing on gravid females may provide effective population suppression while minimizing the emergence of resistance and limiting environmental and non-target impacts (5, 6). Oviposition site selection by gravid females plays a pivotal role in ensuring the survival and proper development of their progeny (7, 8). Previous studies have identified temperature, pH, salinity, nutrient availability, and other physicochemical parameters as key drivers of mosquito larval growth (9, 10). Additionally, a heterogeneous composition of bacterial taxa in breeding habitats constitute a key ecological determinant of mosquito larval environments (10), providing essential nutritional resources for immature stages (11). The composition of breeding site microbiota varies with mosquito species, nature of aquatic habitat, and geographic region (10, 12). Moreover, accumulating evidence indicates that olfactory and visual cues associated with breeding sites act synergistically to shape mosquito oviposition preference (13–15). Volatile infochemicals released by bacteria, soil, plants, predators, and conspecific eggs and larvae shape mosquito sensory perception and guide oviposition site selection, thereby enhancing reproductive success (14, 16–19). Among these cues, microorganisms inhabiting the breeding sites play a pivotal role by producing microbial volatile organic compounds (mVOCs) that modulate mosquito oviposition behavior (16, 20–24). Studies by *Lindh et al*. identified 13 putative oviposition-attracting semiochemicals emitted by the bacterial isolates including aromatic and aliphatic alcohols, aliphatic ketones, alkylpyrazines, dimethyl oligosulfides, and indole (16). Additionally, geosmin, the most common microbial smell of wet land, promote egg-laying site selection in *Aedes aegypti* (20). Recently, 2-ethylhexanol and 2,4-di-tert-butylphenol, produced by *Klebsiella* spp., have been identified as oviposition attractants for *Aedes aegypti* (25). Therefore, elucidating the mechanisms by which this mosquito exploits nutrient-poor artificial breeding habitats could provide critical insights into habitat ecology and facilitate the development of more effective and sustainable vector control strategies.

Pune, a rapidly growing metropolitan city in Maharashtra, in the western region of India, has witnessed repeated sporadic outbreaks of arboviral diseases such as dengue, Zika, and chikungunya since 1970 (26). In the absence of widely effective vaccines, vector control strategies, including insecticide-based interventions such as indoor residual spraying, remain the cornerstone of disease prevention (27). However, the efficacy of commonly used insecticides, including DDT, permethrin, deltamethrin, and lambda-cyhalothrin, is increasingly compromised due to the emergence of resistance in *Ae. aegypti* populations across India (28). Moreover, recent major outbreaks of dengue, chikungunya, and Zika over the past five years also reflects the increasing abundance of mosquito breeding habitats in Pune (26, 29). Therefore, this study aimed to identify microbial signature of urban *Aedes* breeding habitats and compare their microbial diversity with natural habitats colonized by other culicine mosquitoes, including *Culex sp*. Water samples were collected during entomological surveillance conducted during Zika and chikungunya outbreaks in 2024 and subjected to metagenomic analysis. Our results revealed significantly greater bacterial diversity in artificial breeding habitats than in natural habitats, with a marked enrichment of *Enterobacter* sp. in artificial habitats. In contrast, *Hydrogenophaga* and *Methyloparacoccus* were predominant in natural freshwater habitats. Dual-choice oviposition assays demonstrated that *Enterobacter* sp. significantly increased oviposition preference in gravid *Ae. aegypti* females. Whereas, natural habitat associated *Hydrogenophaga* sp. significantly suppressed egg laying relative to the control. Furthermore, the GC-MS profiling of bacterial metabolites identified indole as a putative metabolic fingerprint associated with *Enterobacter* sp. The co-occurrence of *Enterobacter* enrichment, indole production, and enhanced oviposition preference suggests that indole may contribute to the chemical cues underlying breeding-site selection by *Ae. aegypti*.

## Materials and Methods

The technical overview of the current investigation is illustrated in **Figure 1**.

**Figure 1:**
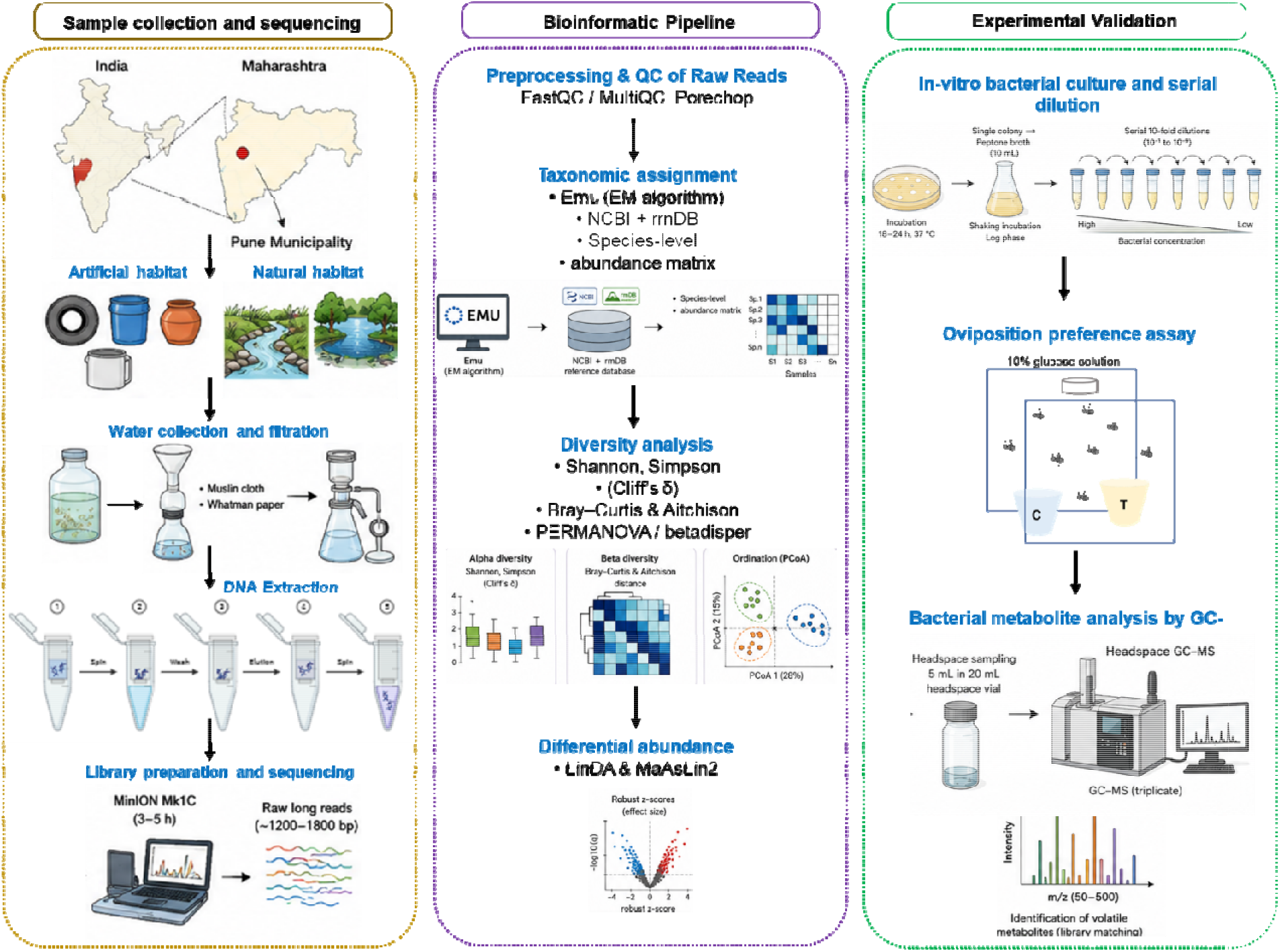
Schematic overview of the study workflow, including sample collection, metagenomic sequencing, bioinformatic analysis and taxonomic assignment, followed by in vitro bacterial culture, oviposition preference assays, and metabolomic profiling of bacterial metabolites for experimental validation.

### Study site and study design

Pune, in western India, has a tropical climate with a mean annual temperature of 24.3°C, average annual rainfall of about 1,200 mm, and mean relative humidity of 69%. The city is traversed by two major rivers, the Mula and Mutha. The study sites were selected based on documented incidences of Zika and chikungunya in 2024, ensuring epidemiological relevance (**Figure 2**). Sampling containers were chosen randomly to minimize selection bias. Water storage containers containing immature stages (larvae or pupae) of *Ae. aegypti* were classified as positive, whereas containers devoid of any developmental stage of *Ae. aegypti* were designated as larval-negative habitat. Given the preference of *Ae. aegypti* for artificial containers and the predominance of *Culex* spp. in natural aquatic habitats, river and lake samples were included to compare bacterial communities across breeding habitats. In total, 24 water samples were subjected to downstream metagenomic analysis to enable comparative profiling of microbial communities across habitat types: 19 from artificial breeding containers, including one larva-negative artificial habitat (FBA13, a container without detected mosquito larvae, used as a reference), and five from natural aquatic habitats.

**Figure 2:**
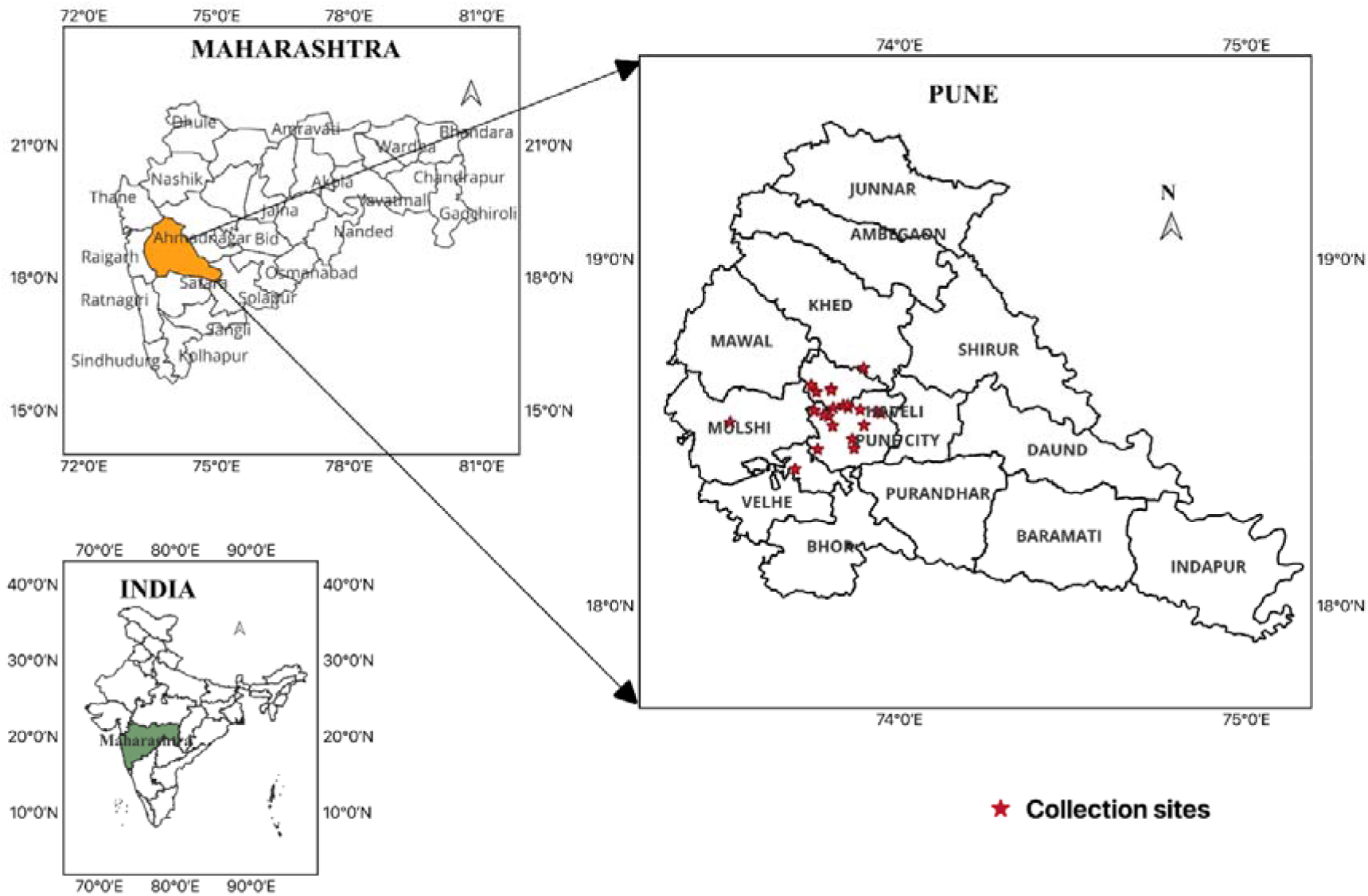
Map of Pune Municipal Corporation highlighting the locations of sample collections (red stars). It is created using QGIS Version 3.44.12

### Collection of water samples and bacterial DNA extraction

Water samples were collected in sterile 500 mL glass bottles using the standard dipping method. Within 24hrs of sampling, the water samples were pre-filtered sequentially through muslin cloth and Whatman filter paper to remove particulate debris. For bacterial enrichment, the filtrate was passed through 0.22 µm membrane filters using a vacuum filtration unit to capture microbial cells. The membranes were aseptically transferred to sterile Petri dishes and stored at −20 °C until further processing.

For DNA extraction, membranes were sectioned into small pieces and incubated in 1.5 mL lysis buffer (10mM Tris-HCl of pH 8.0, 10mM EDTA, disodium salt of pH 8.0, 2% SDS (v/v), 100mM NaCl, 1mg/ml Lysozyme) for 1 hr at 200 rpm in an orbital shaker incubator. Genomic DNA was isolated using the QIAamp DNA Mini Kit (Cat. no. 51304, Qiagen, Hilden, Germany) following the manufacturer’s instructions. Briefly, 500 µL of lysate was mixed with 200 µL of Buffer AL and incubated at 72 °C for 10 min, then 200 µL of ethanol was added. The mixture was loaded onto a spin column and centrifuged at 8000 rpm for 1 min, and the flow-through was discarded. After wash steps, DNA was eluted in 40 µL of pre-warmed nuclease-free water. DNA concentration and purity were assessed using a Nanodrop spectrophotometer (Thermo Fisher Scientific ND-1000), and samples were stored at −20 °C for subsequent analysis. The presence of bacterial DNA in the samples was evaluated by conventional PCR amplification of the conserved 16S rRNA gene region using PCR conditions (95 C for 5 mins, 95 C for 15 sec, 50 C for 30 sec, 72 C for 50 Sec, and final extension at 72 C for 5 mins) (Forward primer: 5’AGAGTTTGATCCTGGCTCAG 3’, reverse primer: 5’GGTTACCTTGTTACGACTT 3’) (30).

### Bacterial full-length 16S rRNA sequence amplification

The concentration of extracted DNA was quantified using the Qubit dsDNA HS Assay Kit with a Qubit 4.0 Fluorometer (Thermo Fisher Scientific, USA) following the manufacturer’s instructions. Multiplexed full-length 16S rRNA gene sequencing libraries were prepared using the SQK-16S024 kit (Cat. No. 16S1424; Oxford Nanopore Technologies, UK). Briefly, 10 ng of DNA template (10 µL) was used for amplification of the full-length 16S rRNA gene (V1–V9 regions) with barcoded primers (primers: 5’-barcode-27F-AGAGTTTGATCMTGGCTCAG-3’, 5’-barcode-1492R-CGGTTACC TTGTTACGACTT-3’) in a reaction containing 25 µL of LongAmp Hot Start Taq 2× Master Mix (Cat. No. M0533L; New England Biolabs, USA) and 5 µL of nuclease-free water. PCR amplification was performed under the following conditions: initial denaturation at 95 °C for 1 min; 35 cycles of 95 °C for 20 sec, 55 °C for 30 sec, and 65 °C for 2 min; followed by a final extension at 65 °C for 5 min. PCR amplicons were purified using 1× AMPure XP beads (Oxford Nanopore Technologies, UK), and DNA concentration was re-quantified using the Qubit 4.0 Fluorometer. Purified amplicons from each sample were pooled in equimolar concentrations to achieve a final library of 60–70 fmol in 10 µL of 10 mM Tris-HCl (pH 8.0) with 50 mM NaCl. Subsequently, 1 µL of rapid adapter was added to the pooled library, gently mixed, briefly centrifuged, and incubated at room temperature for 5 min. The prepared library was then loaded onto an R10 flow cell (Oxford Nanopore Technologies, UK) and sequenced on the MinION Mk1C platform for 3–5 hrs, depending on the number of samples.

### Pre-processing of 16S nanopore amplicon sequencing data

Full-length 16S rRNA gene sequences (V1–V9 regions, ∼1200–1800 bp) were pre-processed by performing extensive quality control followed by taxonomic assignment (**Figure 1**). Initial quality assessment using FastQC (v0.12.1) and MultiQC (v1.32) revealed that most samples exhibited acceptable quality metrics for nanopore long-read data, with Phred scores predominantly greater than 10 across the 1000–2000 bp range. GC content distribution was unimodal, peaking at 52–55%, consistent with bacterial 16S rRNA gene composition. Adapter trimming was performed with Porechop (v0.2.4) to remove barcodes and adaptors from 95% reads and filtering out approximately 200 chimeric reads per sample. Further, the reads were filtered using fastplong (v0.4.1) to retain sequences within the 1200–1800 bp range, with at least 50% of bases having Q-score >10 and a mean Q-score >15. This filtering step removed 3,000–5,000 low-quality or off-target reads per sample while retaining 15,000 to >100,000 high-quality reads, with most samples exceeding 20,000 reads.

Taxonomic assignment was conducted using Emu v3.5.5 (Expectation-Maximization for Unbiased Taxonomic Annotation), which employs compositional abundance estimation algorithm to estimate microbial community composition from error-prone long-read 16S rRNA sequencing data (31). Reads were aligned against the default NCBI+rrnDB reference database. Taxonomic abundance estimates generated at the taxon ID, species, genus, and family levels were consolidated into a species-level relative abundance matrix for subsequent analyses. Species-level assignments based on full-length 16S rRNA sequences, while more resolved than short-read 16S data, do not always provide reliable discrimination among closely related species within some genera. Accordingly, species-level names reported here should therefore be interpreted as Emu-based taxonomic assignments, however, the key conclusions are drawn primarily at the genus level.

### Alpha & Beta diversity analysis

Alpha and beta-diversity analyses comparing natural and artificial samples were conducted using the scikit-bio package (v0.7.1) (32) in Python (v3.12.12). Diversity metrics were visualized using the seaborn (v0.13.2) and matplotlib (v3.10.8) libraries. For beta-diversity analyses, Bray–Curtis and Aitchison distance matrices were computed, and Principal Coordinates Analysis (PCoA) was performed to visualize patterns of sample clustering in reduced-dimensional space. Statistical differences in community composition were assessed using Permutational Multivariate Analysis of Variance (PERMANOVA) (33). To verify the assumption of homogeneity of group dispersions underlying PERMANOVA, tests of multivariate homogeneity of dispersions (betadisper) were subsequently performed (33). Both PERMANOVA and betadisper analyses were implemented using the vegan package (v2.7-3) in R (v4.5.2).

To evaluate the relative contributions of habitat type and sequencing batch to variation in microbial community composition, a PERMANOVA model was fitted using Bray–Curtis dissimilarities. The combined model indicated 18.4% of the variation in community composition (*P* = 0.002). Habitat type accounted for a larger proportion of the variance (11.0%, *P* = 0.026) than flow cell (7.1%, *P* = 0.005). Together, habitat type and flow cell accounted for less than one-fifth of the total variation in community composition, leaving most of the variance unexplained by either factor. However, because all-natural samples were sequenced on a single flow cell (FAW), batch correction was not performed, as it could obscure genuine habitat-associated variation.

### Differential abundance analysis

To discover specific microbial features characterizing natural versus artificial habitats, we performed differential abundance analysis to identify taxa whose enrichment was associated with habitat type. Given the modest sample size (n=24) and pronounced class imbalance (5 natural vs. 19 artificial habitats), we used a consensus-based approach integrating two complementary differential-abundance methods: LinDA (Linear models for Differential Abundance analysis of microbiome compositional data) (34) and MaAsLin2 (Microbiome Multivariable Associations with Linear Models) (35), both applied to Emu’s native estimated-count species table (uncorrected for sequencing batch). Because all five natural-habitat samples were sequenced on a single flow cell (FAW) while artificial-habitat samples spanned two flow cells (FAW and FBA), habitat type and sequencing batch are structurally confounded in the full dataset. This confound cannot be fully disentangled statistically within the complete dataset, and is acknowledged as an inherent limitation of the study design. To address it as far as possible, flow cell was included as a covariate in the primary differential-abundance model (habitat type + flow cell; *n* = 23 biological samples), excluding the larva-negative artificial-habitat sample FBA13. As an additional sensitivity analysis, the same consensus differential-abundance analysis was repeated using only samples sequenced on the FAW flow cell, comprising 12 artificial-habitat and 5 natural-habitat samples (*n* = 17). Restricting the analysis to a single flow cell minimized potential confounding by inter-flow-cell sequencing batch effects, thereby providing an independent validation of habitat-associated differential-abundance patterns. Species were retained for differential-abundance testing only if they met a stringent prespecified filtering criterion (prevalence ≥20% of samples, mean relative abundance ≥0.1%, and absence from the larva-negative artificial-habitat sample FBA13). This filtering procedure was applied independently to each of the two sample sets described above. A species was designated as a primary candidate habitat-associated taxon only if it achieved statistical significance (*q* < 0.05, Benjamini–Hochberg adjusted) in both LinDA and MaAsLin2 analyses, and in both the flow-cell-adjusted analysis of the full dataset and the single-flow-cell replication analysis. Thus, primary candidate taxa were restricted to those showing consistent habitat associations across both statistical frameworks and both approaches to addressing the potential sequencing-batch confound.

### Metabolic pathway prediction

To further explore potential mechanistic links between habitat-associated bacterial communities and mosquito oviposition behavior, we used PICRUSt2 (Phylogenetic Investigation of Communities by Reconstruction of Unobserved States 2) (36) to predict community-wide MetaCyc metabolic pathway abundances from the full 16S rRNA taxonomic profile (1,221 species; stratified, per-sequence contribution mode enabled). Pathways detectable in the larva-negative artificial habitat (FBA13) were excluded to reduce likely background/contaminant signal. Predicted pathway abundances were tested for differential abundance between habitats using LinDA (34), and pathways significant at FDR q≤0.05 (Benjamini-Hochberg) were retained. PICRUSt2 functional predictions are inferred from 16S taxonomic composition rather than measured directly, and because a significant pathway-level difference does not by itself identify which community members drive it or demonstrate a specific metabolic mechanism. Therefore, these findings are interpreted as hypothesis-generating rather than confirmatory. To relate significant pathways to the taxa identified above, PICRUSt2’s stratified per-taxon contribution profiles were examined for each significantly different pathway. This analysis identified which primary candidate habitat-associated species contributed to each pathway and quantified their relative contributions to the predicted pathway abundance.

### In-vitro bacterial culture

To investigate the association of bacterial genera with mosquito behavior, representative species of the genera *Enterobacter* and *Aeromonas (Enterobacter cloacae* and *Aeromonas caviae)* were revived from glycerol stocks (−80 °C) and cultured on Luria-Bertani (LB) agar (HiMedia Laboratories, Mumbai, India) plates using the quadrant streaking technique, followed by aerobic incubation at 37 °C for 18–24 hrs. In contrast, *Hydrogenophaga crocea* BA0156 (= MCC 3062 ; GenBank CP049989; GCF_011388215.1) (37) was cultured on Luria-Bertani (LB) agar (HiMedia Laboratories, Mumbai, India), supplemented with 0.5% NaCl (pH 7.0) and incubated at 30 °C for up to 5 days to accommodate their slower growth kinetics. Well-isolated single colonies were subsequently inoculated into 10 mL sterile peptone broth (HiMedia Laboratories, Mumbai, India) and incubated under shaking conditions to obtain log-phase cultures. Bacterial cell density was quantified by performing a series of 10-fold serial dilutions (10 ¹ to 10) in sterile phosphate-buffered saline (PBS). Aliquots from these dilutions were plated, and colony-forming units (CFU/mL) were calculated to determine viable bacterial counts and standardize inoculum concentrations for downstream assays using the formula Colony forming unit/ml (CFU/mL) = (No. of Colonies × Dilution Factor) / (Volume of suspension inoculate × Volume of Broth)

### Oviposition Preference Assay

To determine the influence of bacterial-derived cues likely influencing oviposition behavior, a dual-choice oviposition assay was conducted using a laboratory-reared colony of *Ae. aegypti*. Adult females (6–7 days post-emergence) were blood-fed on live chicken feet for approximately 1 hr. Fully engorged females were separated and maintained under standard insectary conditions with access to a glucose source for 72 hrs. to facilitate oocyte maturation. For the assay, cohorts of 10 gravid females were introduced into experimental cages containing two oviposition cups arranged in a randomized spatial configuration: a test cup containing a bacterial suspension (2-4 × 10 CFU/mL) and a control cup containing sterile phosphate-buffered saline (PBS), each with a final volume of 30 mL. Whatman filter paper was provided in each cup as an oviposition substrate. After a 24 hrs of exposure period, the filter papers were retrieved, and the number of eggs laid in each treatment was quantified to assess oviposition preference. The difference in egg deposition between the bacterial treatment and control was evaluated using a paired *t*-test and Wilcoxon Signed-Rank test.

### Bacterial metabolite Analysis

Fresh cultures of *Enterobacter cloacae* and *Aeromonas caviae* were grown in 5ml of sterile peptone water at a dilution of 2-4×10 CFU/mL and centrifuged at 6,000 rpm for 15 minutes at 4 °C to pellet the cells. The resulting cell pellet was washed three times and resuspended in 5 mL of sterile peptone water. The resuspended cell pellets of *Enterobacter cloacae* and *Aeromonas caviae* were subjected to headspace GC–MS analysis. Sterile peptone water served as the control. For each sample and control, 5 mL was transferred into 20 mL headspace vials, which were then securely sealed prior to analysis.

Headspace volatile compounds were analyzed using a Shimadzu GCMS-TQ40 gas chromatograph coupled to a triple quadrupole mass spectrometer (Shimadzu Corporation, Kyoto, Japan). Chromatographic separation was achieved on a TG-5MS fused-silica capillary column (30m × 0.25mm i.d., 0.25μm film thickness; Thermo Scientific, USA). Helium (99.999% purity) was used as the carrier gas at a constant flow rate of 2.38 mL min ¹. Samples were introduced in splitless injection mode, and the injector temperature was maintained at 250 °C. The oven temperature program was as follows: an initial temperature of 60 °C, held for 1 min, ramped at 25 °C min ¹ to 300 °C, and held for 2.5 min, resulting in a total run time of 13.10 min. The mass spectrometer was operated in electron ionization (EI) mode at 70 eV. The ion source and interface (transfer line) temperatures were maintained at 250 °C and 300 °C, respectively. Mass spectra were acquired in full-scan mode over an m/z range of 50–500. Each sample was analyzed in triplicate. The GC-MS data were first cleaned by excluding volatiles present in both the blank and the peptone water control. Volatile compounds were identified and quantified by comparing the acquired mass spectra and retention time with those in the NIST Mass Spectral Library (Version 2.2).

## Results

### Metagenomic profiling and community composition

A total of 24 water samples were collected from multiple locations across the Pune metropolitan region during the 2024 Zika and chikungunya outbreak (**Table S1**; **Figure 2**). All samples were subjected to 16S rRNA amplicon sequencing using the Oxford Nanopore MinION platform to enable comprehensive characterization of the associated microbial communities. Following quality filtering and taxonomic classification using Emu (31), a total of 1,221 bacterial species representing 509 genera were identified. A total of 23 bacterial phyla were identified, with Proteobacteria consistently dominating the microbial communities across most samples and exhibiting the highest relative abundance in both habitat types (**Figure 3A**). In addition to Proteobacteria, Bacteroidetes, Firmicutes, Planctomycetes, and Actinobacteria were among the predominant phyla, with substantial variation in their relative abundances across samples. Microbial community composition displayed pronounced inter-sample variability, with artificial habitats exhibiting greater taxonomic heterogeneity relative to natural habitats. At the genus level, the 10 most abundant genera included *Acidovorax*, *Enterobacter*, *Curvibacter*, *Rhodobacter*, *Hydrogenophaga*, *Aeromonas*, *Cnuella*, *Limnohabitans*, *Polynucleobacter*, and *Lacisediminimonas* (**Figure 3B).**

**Figure 3:**
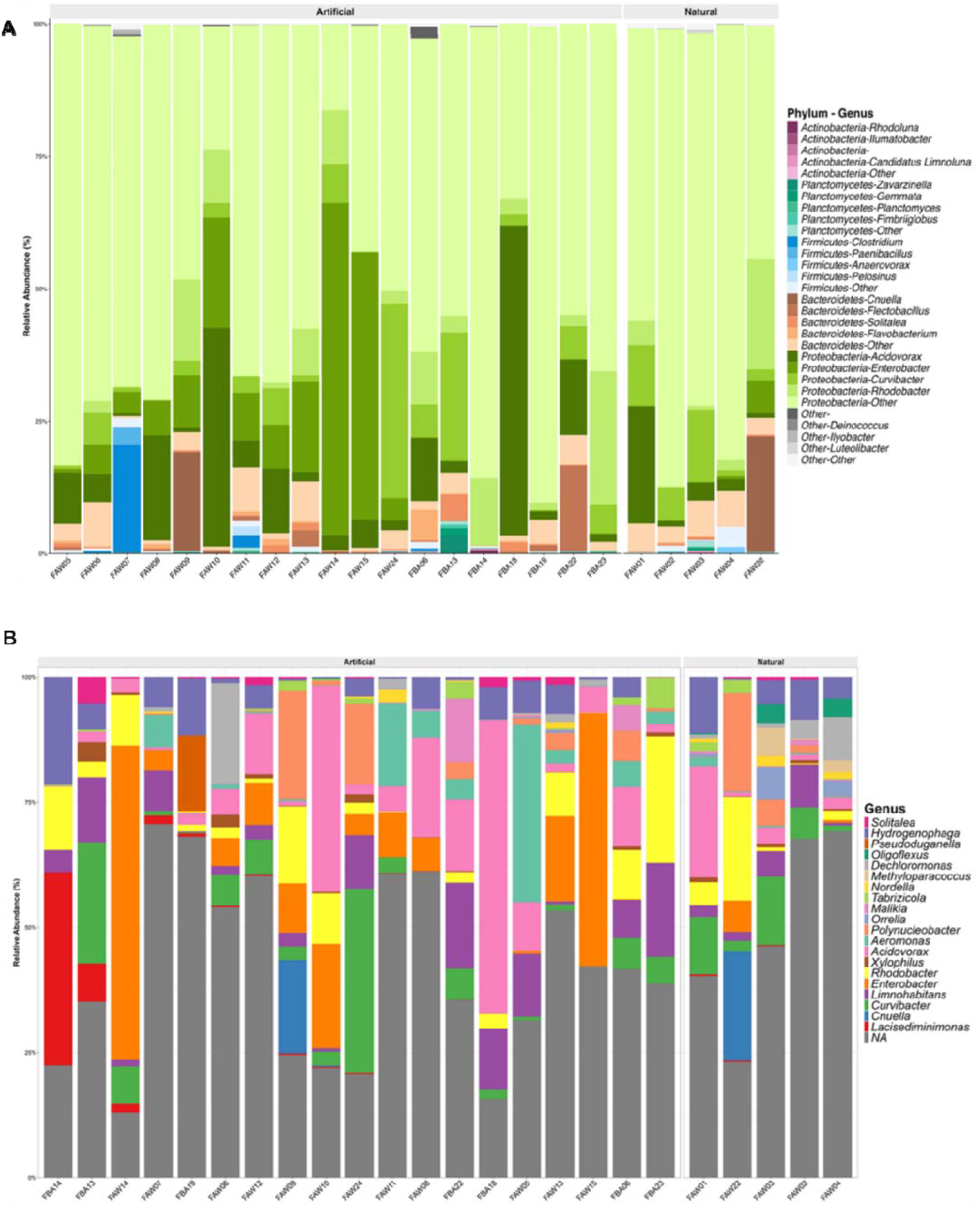
The microbiome composition of artificial and natural habitats. (A) Phylum-level and (B) genus-level compositional difference and relative abundance per sample (top 20 genera; remaining taxa pooled as ‘Other’/‘NA’), according to habitat type (artificial vs natural). N = 24 (19 Artificial, including the larva-negative habitat FBA13; 5 Natural).

Notably, *Enterobacter* was highly enriched in several artificial habitat samples but was largely absent from natural water bodies. Similarly, *Acidovorax* and *Aeromonas* exhibited elevated relative abundances across multiple artificial habitats. In contrast, *Hydrogenophaga* and *Methyloparacoccus* were more prevalent in natural freshwater systems, whereas *Rhodobacter* exhibited no clear habitat preference and showed a heterogeneous distribution across both habitat types (**Figure 3B)**. The larva-negative artificial habitat sample (FBA13) exhibited a distinct microbial signature characterized by the highest relative abundance of the phylum Planctomycetes and the genus *Solitalea* relative to all other samples (**Figure 3A, B**). Its taxonomic composition resembled an intermediate state between natural and artificial habitats and lacked the characteristic enrichment of artificial habitat-associated genera, particularly *Enterobacter* and *Aeromonas*.

### Community diversity across habitat types

To assess within-sample species richness and evenness across different habitat types, alpha-diversity was estimated using the Shannon and Simpson diversity indices. Since Emu provides species-level relative abundance profiles for each sample, rarefaction-based normalization to correct for sequencing depth variation was not applied, thereby minimizing potential biases associated with subsampling procedures (31). Sequencing depth varied considerably among samples (range: 13,220–105,420 reads; **Table S2**). To assess whether differences in sequencing depth influenced the alpha-diversity comparisons, samples were additionally rarefied to the minimum observed sequencing depth, and Shannon and Simpson diversity indices were recalculated. The rarefied analysis yielded results materially consistent with those obtained from the unrarefied data reported below. This consistency, together with the observation that the per-sample rarefaction curves had plateaued before reaching the minimum observed sequencing depth (**Figure S1, Table S3**), indicates that sequencing depth did not substantially influence the observed alpha-diversity patterns. Artificial habitats exhibited significantly greater alpha-diversity than natural water bodies, as reflected by both the Shannon index (median: 3.61 vs. 2.92; Wilcoxon rank-sum test, *P* = 0.019) (**Figure 4A)** and the Simpson index (median: 0.94 vs. 0.85; *P* = 0.003) (**Figure 4B**) **(Table 1**). Despite the limited sample size and substantial variability among natural habitats (n = 5), effect size estimation using Cliff’s delta (38) revealed large effects for both diversity metrics (Shannon: δ = 0.68, 95% CI = 0.23–0.89; Simpson: δ = 0.83, 95% CI = 0.47–0.95) (**Table 1**), indicating strong and consistent enrichment of alpha-diversity in artificial habitats. The larva-negative artificial habitat sample (FBA13) showed substantially reduced alpha diversity relative to the majority of artificial habitat samples, with diversity values falling at or below the first quartile of the artificial habitat distribution (**Figure 4A, B**). Notably, its diversity metrics were comparable to those observed in natural habitats (**Figure 4A, B**). Overall, the observed differences in microbial diversity likely reflect the contrasting environmental characteristics of artificial and natural aquatic habitats (9).

**Figure 4.**
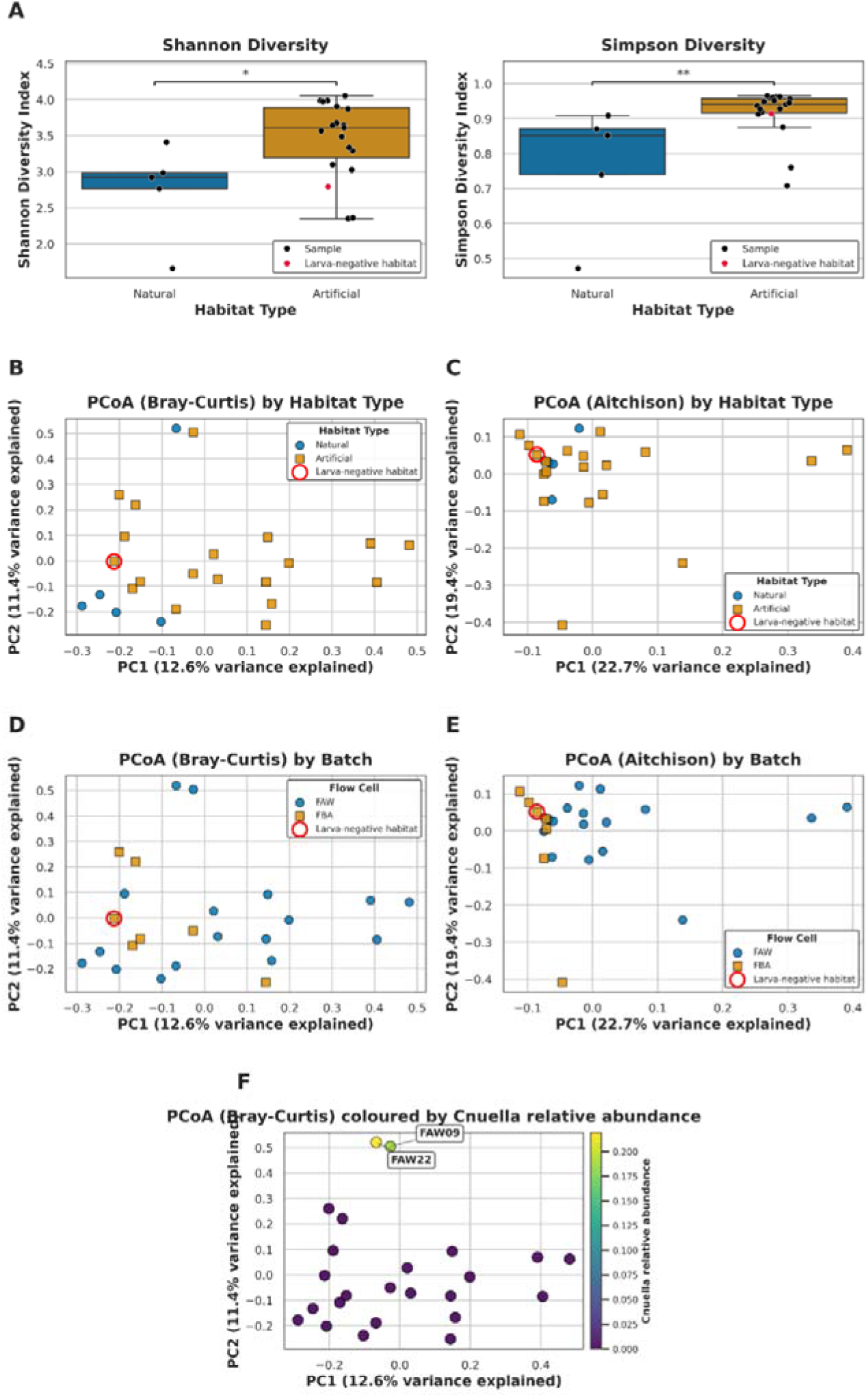
Diversity profiles of mosquitoes’ microbiome. (A) Alpha diversity: box plots of Shannon and Simpson indices for natural versus artificial habitats (points, individual samples; larva-negative artificial habitat FBA13 in red circle). (B–F) Beta diversity: principal-coordinates analysis (PCoA) of (B) Bray-Curtis and (C) Aitchison distances coloured by habitat type, and (D) Bray–Curtis and (E) Aitchison distances coloured by sequencing batch/flow cell (FAW/FBA); (F) Bray-Curtis PCoA coloured by *Cnuella* relative abundance, highlighting the outliers FAW22 and FAW09. Significance: *P<0.05, **P<0.01.

**Table 1:**
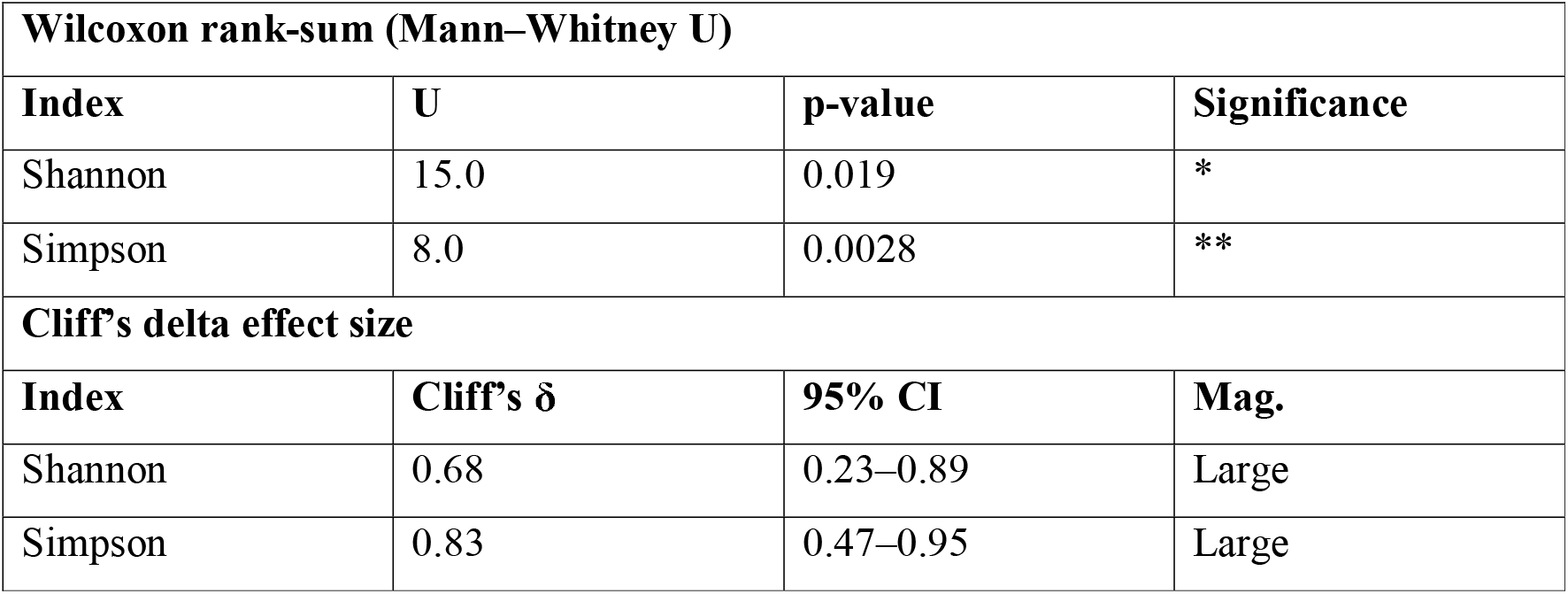
Statistical tests of community diversity analysis.

Next, beta-diversity analysis, based on Bray-Curtis and Aitchison distances, was performed to evaluate differences in bacterial community composition among samples. Principal coordinate analysis (PCoA) based on both metrics showed substantial overlap between natural and artificial habitats, with no clear clustering by habitat type (**Figure 4B–E**). The larva-negative artificial habitat sample (FBA13) also clustered with the remaining samples, suggesting that overall community structure was broadly similar across habitats (**Figure 4B– E**). These results indicate that mosquito attraction and oviposition are likely associated with specific bacterial taxa rather than large-scale shifts in community composition. Notably, two samples (FAW22 and FAW09) appeared as outliers in the Bray-Curtis ordination, likely driven by their high relative abundance of *Cnuella* (**Figure 4F**).

### Differential Abundance Analysis: Identifying habitat-associated bacterial taxa

Analysis of the species-level abundance matrix using LinDA and MaAsLin2, applying a stringent combined filter (prevalence ≥20% of samples, mean relative abundance ≥0.1%, and absence from the larva-negative artificial habitat FBA13), identified 19 species significant (q≤0.05) by LinDA and 15 by MaAsLin2 in the flow-cell-adjusted model (habitat_type + flow_cell, n=23), out of the 71 species passing the filter. Of these, 15 species were consistently identified by both methods (13 Natural-enriched, 2 Artificial-enriched). The two methods showed high concordance in effect estimates, with a Spearman’s rank correlation of ρ = 0.98, a Lin’s concordance correlation coefficient of 0.98, and 90% agreement in the direction of association (**Figure S2**). To determine which of these associations could be attributed to sequencing batch rather than habitat, the same consensus analysis was independently repeated using only the single flow cell (FAW) on which both habitat types are represented (n=17 samples). The filtering criteria yielded 78 taxa eligible for testing, which by construction cannot be confounded by flow cell. Thirteen of the 15 taxa identified in the full dataset remained significant in both LinDA and MaAsLin2 (q≤0.05) in this single-flow-cell analysis (including 11 Natural-habitat-enriched, 2 Artificial-habitat-enriched). These 13 taxa were therefore defined as the primary set of candidate habitat-associated taxa for subsequent analyses (**Figure 5A, B; Table S4**). Of these, taxa belonging to the genera *Arcobacter, Bacteroidales, Methyloparacoccus, Novosphingobium, Rhodobacter, Bordetella, Dechloromonas, Hydrogenophaga, Achromobacter*, and *Quisquiliibacterium* (**Figure 5A, C**) were enriched in natural habitats. In contrast, *Enterobacter* (*E. asburiae* and *E. roggenkampii*) was the only genus identified as significantly enriched in artificial habitats by both analytical methods **(Figure 5A, C)**. No other artificial-habitat-associated taxon met the predefined criteria for inclusion as a primary candidate.

**Figure 5.**
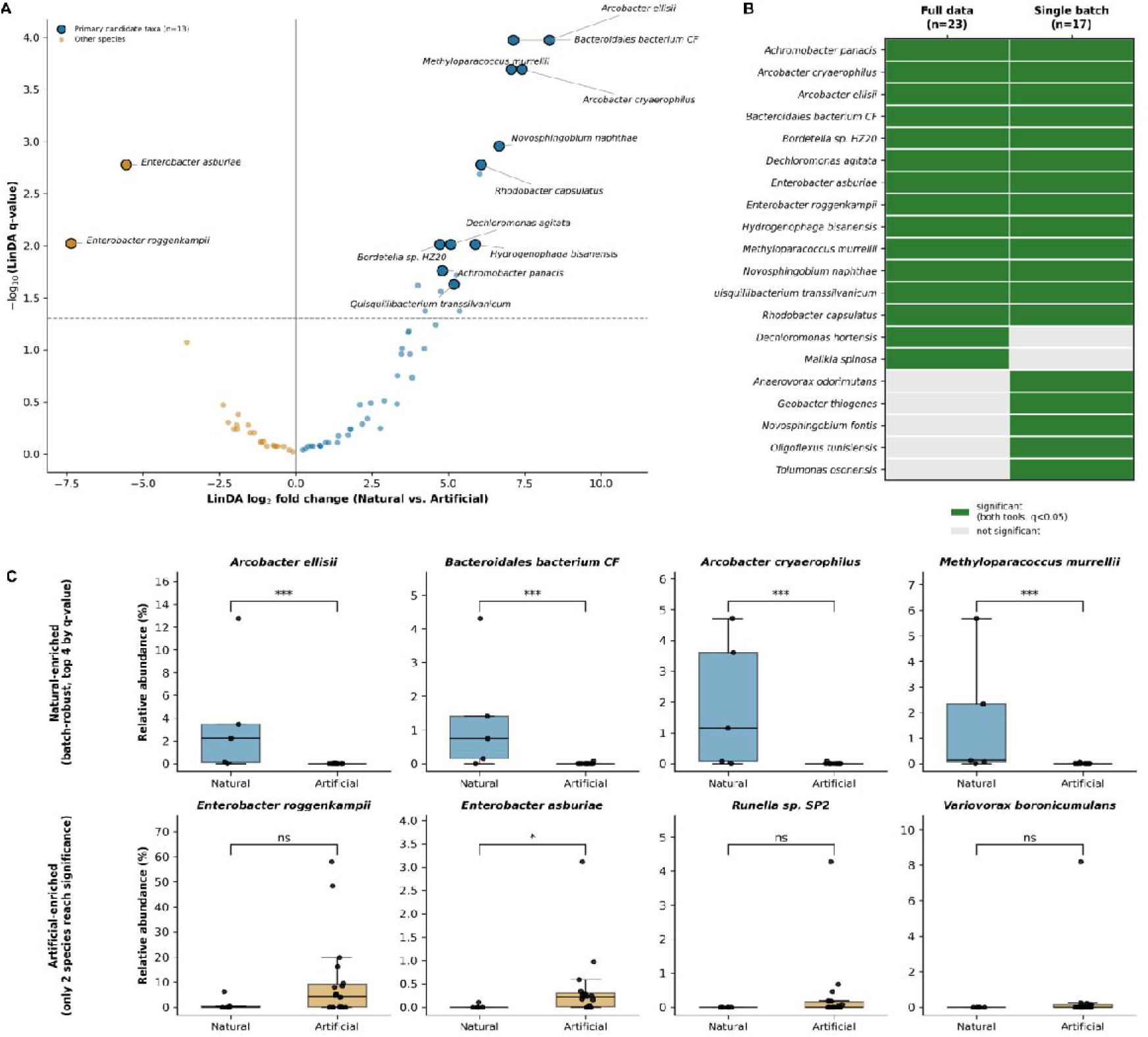
Consensus differential-abundance analysis identifies batch-robust habitat-associated bacterial taxa. (A) Volcano plot of LinDA log fold change vs. q-value for the species passing the stringent filter (habitat_type + flow_cell model, n=23); points are coloured by two categories: the 13 primary candidate taxa (significant, q≤0.05, both LinDA and MaAsLin2, in both the flow-cell-adjusted model and the single-flow-cell replication; labelled) and all other species. (B) Batch-sensitivity heatmap: significance (both tools, q≤0.05) of each candidate taxon in the full dataset (flow-cell-adjusted, n=23) and the single-batch replication (FAW-only, n=17); taxa significant in both form the primary candidate set. (C) Relative abundance (%) of representative taxa in natural versus artificial habitats: the four Natural-enriched primary candidates with the lowest q-values (top row), and the two Artificial-enriched primary candidates (*Enterobacter asburiae, E. roggenkampii*) alongside the two next-largest (non-significant) Artificial-leaning taxa shown for visual comparison (bottom row). Boxes show median and interquartile range; points are individual samples. Asterisks: Wilcoxon rank-sum test on relative abundance (ns, not significant; *P<0.05; ***P<0.001); this simple univariate test is less sensitive than the compositional LinDA/MaAsLin2 models used for formal significance calling in (A) and (B). Thirteen taxa met the primary batch-robust candidate criterion (11 Natural-enriched, 2 Artificial-enriched); see Table S4 for the full candidate-selection criteria

Further, to investigate functional differences between microbial communities associated with natural and artificial breeding habitats, metabolic pathway prediction was performed using the MetaCyc database on the full bacterial community. Seven pathways were significantly enriched in natural habitats after FDR correction (LinDA, q≤0.05): L-isoleucine biosynthesis IV, adenosylcobalamin (vitamin B) biosynthesis I, pyruvate fermentation to propanoate I, CMP-pseudaminate biosynthesis, the super-pathway of UDP-N-acetylglucosamine-derived O-antigen building blocks biosynthesis, protein N-glycosylation (bacterial), and ergothioneine biosynthesis I (**Table S5; Figure S3**). Cross-referencing PICRUSt2’s per-taxon stratified contributions showed that only one of these seven pathways, CMP-pseudaminate biosynthesis, is attributable to a primary candidate habitat-associated species (*Arcobacter cryaerophilus*) (**Table S5**). However, no pathways were significantly enriched in artificial habitats after multiple-testing correction.

### *Enterobacter* influence oviposition preference of *Ae. aegypti*

To investigate whether bacterial taxa enriched in artificial breeding habitats influence oviposition site selection by *Ae. aegypti*, binary-choice oviposition assays (**Figure 6A**) were conducted using in vitro-cultured *Enterobacter cloacae* and *Aeromonas caviae* isolates. Gravid females exhibited a significant preference for oviposition in cups containing *E. cloacae* (2-4 × 10 CFU/mL), depositing substantially more eggs than in the peptone water control (Paired t-test = *p* ≤ 0.005 and Wilcoxon Signed-rank test = *p* ≤ 0.01) (**Figure 6B**). In contrast, no significant difference in oviposition preference was observed between *A. caviae-* treated cups and the control, with gravid females laying comparable numbers of eggs in both treatments (**Figure 6C**).

**Figure 6:**
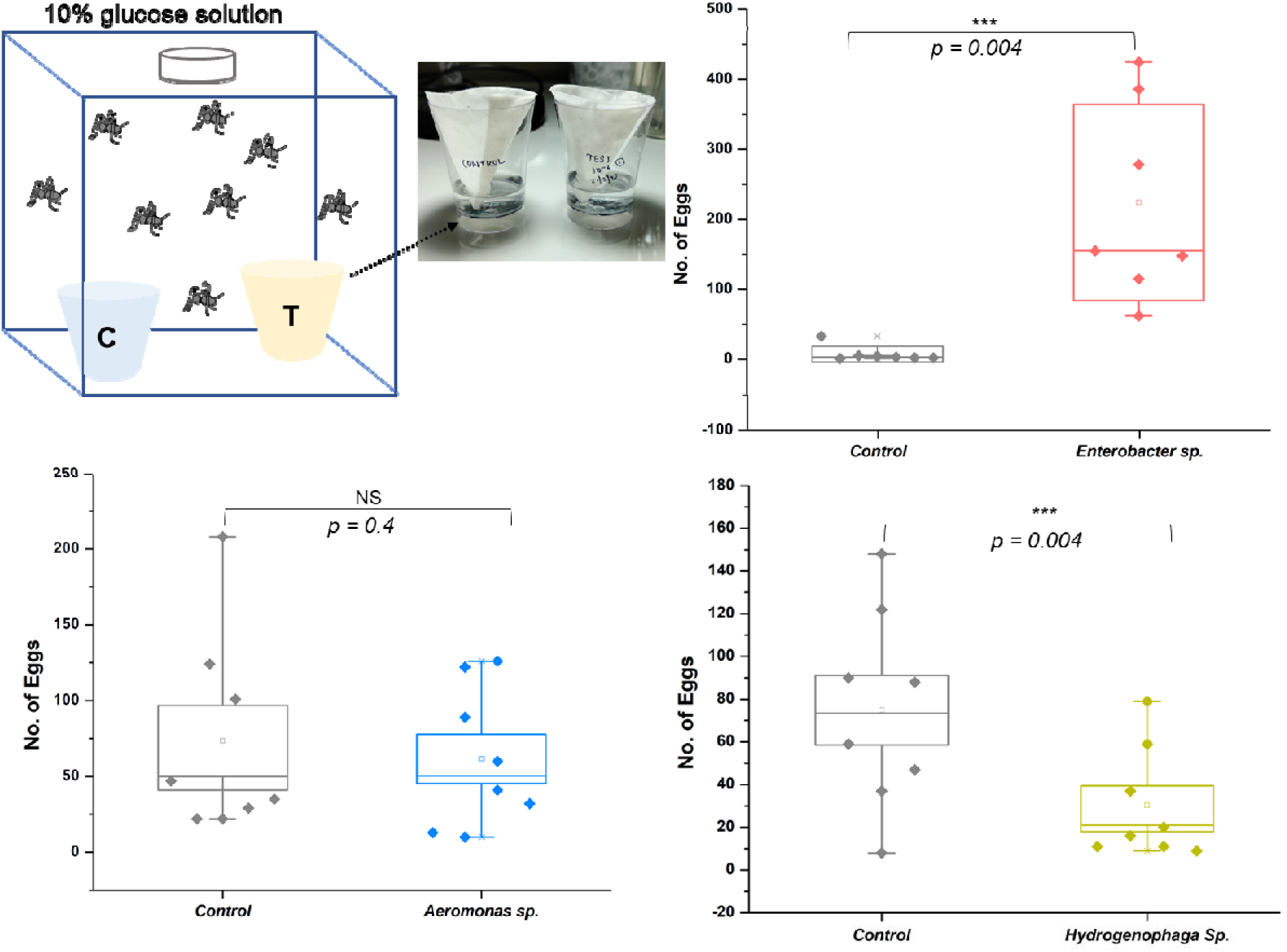
Two-choice laboratory oviposition assays testing oviposition preference for different bacteria. (A) Schematic representation of the dual-choice assay comparing eggs laid in a bacterial test cup (T) versus sterile peptone water control cup (C). (B) Number of eggs laid by gravid females in control vs *Enterobacter* test oviposition cups (p ≤ 0.005). (C-D) Oviposition preference of gravid females between peptone water control cup and test cup containing *Aeromonas* and *Hydrogenophaga* respectively. Each data point represents the number of eggs laid for each biological replicate. The number of mosquitoes used in each assay (n) = 10, and the number of biological replicates (N) = 8 for *Enterobacter* and *Aeromonas* and N = 7 for *Hydrogenophaga*. Values represent the mean ± SEM of 8 and 7 biological replicates. Statistical significance for differences in the number of eggs were computed using the Paired *t*-test and Wilcoxon Signed-rank test. Significance: (*) = p < 0.05, (**) = p < 0.01, (***) = p < 0.001, NS = Not significant

To further evaluate the influence of bacteria associated with natural aquatic habitats, the dominant natural habitat-associated bacterium, *Hydrogenophaga crocea* BA0156, was assessed using the same binary-choice oviposition assay. Gravid females exhibited a significant avoidance of *H. crocea*-supplemented oviposition cups, depositing significantly more eggs in the control peptone-water cups (Paired t-test = *p* ≤ 0.005 and Wilcoxon Signed-rank test = *p* ≤ 0.01) (**Figure 6D**). Taken together, the oviposition preference assays demonstrate that *Enterobacter* sp. promote oviposition site selection, whereas *Hydrogenophaga* sp. exerts an oviposition-deterring effect in gravid *Ae. aegypti* females.

### Bioactive compounds identified from *Enterobacter* headspace

Previous studies have demonstrated that bioactive bacterial metabolites serve as important chemical cues facilitating multiple mosquito behaviors, including host-seeking and oviposition site selection (14, 16, 20). The enhanced oviposition preference of gravid females for *Enterobacter-*treated oviposition cups prompted us to characterize the volatile metabolites associated with bacterial activity that may mediate this behavioral attraction. Accordingly, volatile organic compounds (VOCs) released from actively growing *Enterobacter* cultures were profiled by GC–MS. Parallel profiling of *Aeromonas* cultures enabled comparative assessment of bacteria-specific volatile signatures. After normalization against peptone water controls and blank samples, indole and 2-tridecanone were identified as significantly enriched volatile metabolites in both *Enterobacter* (**Figure 7A);** and *Aeromonas* cultures (**Figure 7B).** Additionally, the alkylphenol 2,4-di-tert-butylphenol was significantly enriched specifically in *Aeromonas* cultures **(Figure 7B)**.

**Figure 7:**
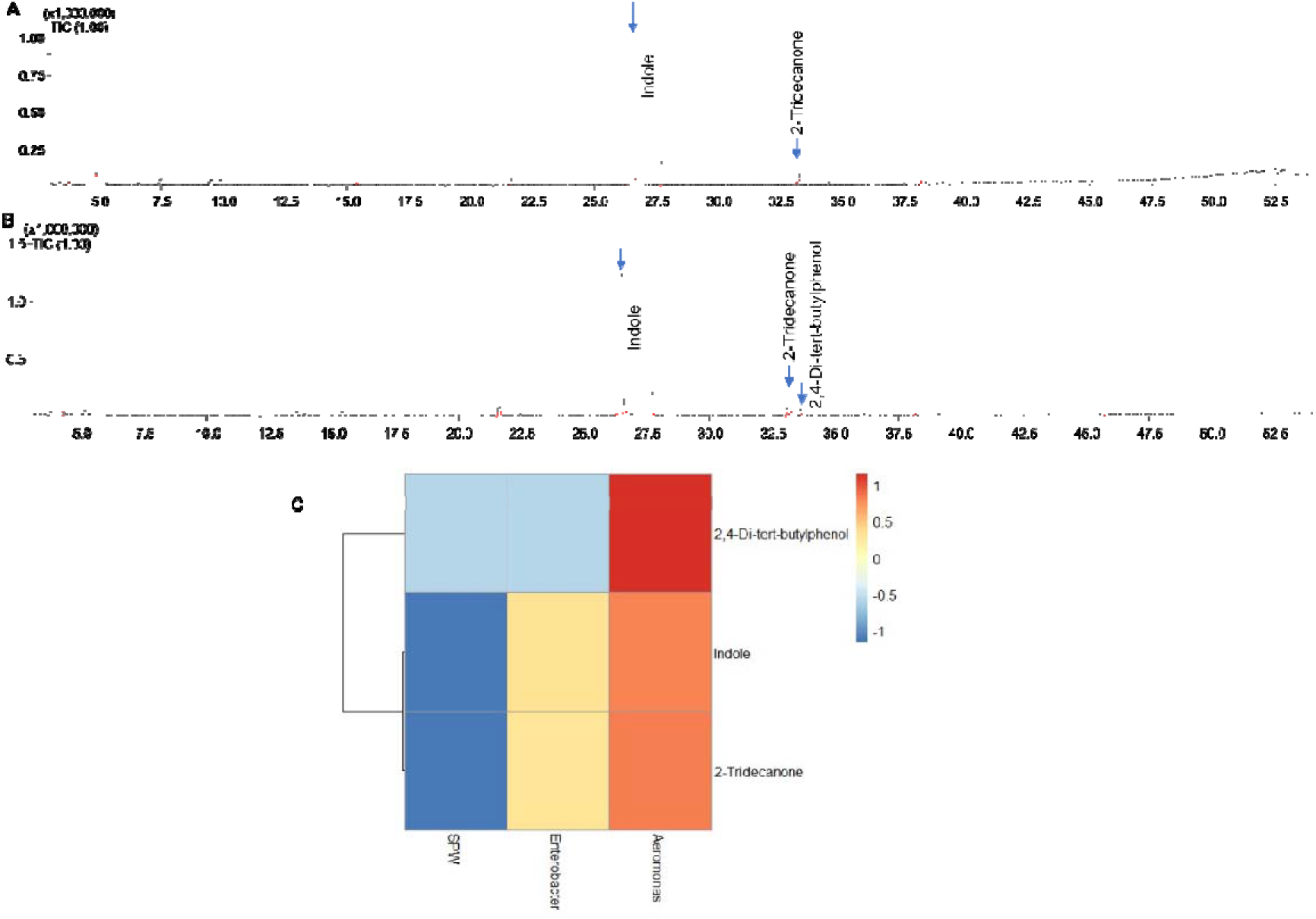
Head-space profiling of volatile bioactive compounds produced by. *Enterobacter* and *Aeromonas* cultures. (A) GC-MS chromatogram of the bacterial metabolites produced by *Enterobacter* cell suspension. (B) GC-MS chromatogram of the bacterial metabolites produced by *Aeromonas* cell suspension. (C) Heatmap depicting the relative abundance of the three predominant VOCs implicated in oviposition site selection across the different sample types. SPW, sterile peptone water; *Enterobacter: Enterobacter* cell suspension; *Aeromonas: Aeromonas* cell suspension

## Discussion

*Ae. aegypti*, the primary vector of several medically important arboviruses, thrives in urban environments by preferentially exploiting anthropogenic water-holding containers for oviposition. Rapid urbanization and inadequate water management have increased the availability of these breeding habitats, contributing to mosquito proliferation and arboviral transmission. Although microbial communities are recognized as modulators of oviposition behavior through the production of semiochemicals, the microbial ecology of *Ae. aegypti* breeding habitats in Indian urban settings remains largely unexplored. Addressing this knowledge gap, the present study integrated habitat-associated bacterial taxa with mosquito oviposition responses and the putative volatile chemical signatures.

### Positive Association of *Enterobacter* with anthropogenic water-holding containers

Despite differences in bacterial diversity and community composition, Proteobacteria dominated both artificial and natural breeding habitats, consistent with previous studies (39). This widespread dominance likely reflects their metabolic versatility, stress tolerance, and ability to persist under nutrient-limited conditions (39). Interestingly, Artificial breeding habitats exhibited significantly higher alpha diversity and were enriched with *Enterobacter*, *Acidovorax*, and *Aeromonas*, whereas natural habitats were characterized by a greater abundance of *Hydrogenophaga* and *Methyloparacoccus.* The observed habitat-specific bacterial communities likely arise from differences in physicochemical conditions and nutrient availability, although these parameters were not measured in the present study. Artificial water-storage containers are exposed to variable organic inputs, human-associated contamination, and fluctuating environmental conditions, creating diverse ecological niches that may promote a broader range of bacterial taxa (9). In contrast, natural aquatic habitats may exhibit specific environmental regimes that select for microbial taxa adapted to these conditions (40, 41). Overall, the bacterial taxa identified in this study are consistent with previous reports from mosquito breeding habitats across diverse geographical regions, where genera including *Pseudomonas*, *Aeromonas*, *Acinetobacter*, and *Enterobacter* have been frequently detected (42–44). In particular, members of the Enterobacteriaceae have repeatedly been associated with oviposition-active breeding sites and mosquito gut microbiota (45–47). Though members of the *Enterobacter* genus are ubiquitous in the environment, their abundance in specific larval habitats suggests a potential ecological association between mosquito habitat selection and anthropogenic/human activities in the surrounding environment. Collectively, these findings highlight the distinct microbial landscape in artificial breeding habitats that may play an important role in shaping mosquito ecology, where *Enterobacter* is likely to impact oviposition behavior significantly.

### *Enterobacter* promotes oviposition-site selection, likely through indole production

The marked enrichment of the genus *Enterobacter* in artificial breeding habitats prompted us to evaluate its effect on oviposition behavior of gravid *Ae. aegypti* females. In a dual-choice oviposition assay, *Enterobacter cloacae*, a laboratory-available representative of the genus, elicited a strong oviposition response, with more than 90% of females exhibiting a preference for the bacterial treatment over the sterile-water control. However, the *Aeromonas* isolate, *Aeromonas caviae*, available in our laboratory, consistent with previous observations, showed no effect on the egg-laying behavior (48). In contrast, the *Hydrogenophaga* isolate, *Hydrogenophaga crocea* BA0156, inhabiting natural water bodies, elicited negative effects and approximately 80% of gravid females exhibiting preference for control cups. Although both *Enterobacter* and *Aeromonas* were enriched in artificial breeding habitats, only *Enterobacter* significantly enhanced oviposition by gravid *Ae. aegypti.* This is consistent with previous reports that *Enterobacter cloacae* and *Bacillus cereus* have been reported to stimulate oviposition in *Aedes aegypti* (5, 49, 50). Moreover, bacterial isolates such as *Aeromonas hydrophila*, *Klebsiella variicola*, *Enterobacter hormaechei*, *Bacillus marisflavi*, and *Aeromonas veronii* exhibited either complete or partial inhibition of oviposition, whereas *Aeromonas taiwanensis* and *Klebsiella* sp. MC1F elicited weak to considerable oviposition attraction (47, 48). These findings suggest that mosquito oviposition behavior may be strongly influenced by the bacterial species present within a given breeding habitat. Similarly, the ecological relevance of the deterrent effect associated with *Hydrogenophaga* may reflect the characteristics of natural breeding habitats, where the presence of substantial industrial and organic waste could generate environmental conditions that are less favorable for oviposition by *Ae. aegypti*.

Previous studies have shown that bacterial communities produce diverse and species-specific volatile chemical signatures, which may contribute to discriminating bacterial taxa. This chemical diversity may influence mosquito oviposition site selection through differential responses to bacterial volatile blends (19–22). Our GC–MS analysis identified several shared volatile metabolites in *Enterobacter* and *Aeromonas,* including indole and 2-tridecanone, both previously implicated in mosquito oviposition behavior (5, 51, 52). However, despite this overlap, only *Enterobacter* elicited a significant oviposition response, indicating that oviposition is determined not by the presence of individual compounds alone but by the composition, relative abundance, and synergistic interactions within the entire volatile blend. The higher abundance of indole and 2-tridecanone in *Aeromonas* cultures may have exceeded attractive thresholds, potentially resulting in a non-significant effect or sensory masking. Moreover, the exclusive detection of 2,4-di-*tert*-butylphenol in *Aeromonas* suggests that certain bacterial metabolites may antagonize otherwise attractive odor blends. Notably, geosmin, a microbial volatile previously reported as an oviposition attractant (20), was not associated with the bacterial communities identified here, suggesting that indole-associated signaling may represent a more ecologically relevant oviposition cue in urban artificial breeding habitats in India. This finding raises the possibility of vector convergence in shared urban habitats, where *Enterobacter*-associated indole signaling may influence both *Ae. aegypti* and the urban malaria vector *An. stephensi* (53). Given the increasing geographic expansion and invasive potential of *An. stephensi* (54), shared microbial cues could promote habitat overlap between these vectors and potentially influence malaria transmission dynamics in tropical urban environments.

## Limitations

Microbial community composition can vary substantially across spatial and temporal scales. In the present study, sampling was geographically restricted to selected sites within the Pune metropolitan region and temporally limited to the winter-to-pre-monsoon period. Therefore, the microbial communities identified may not fully represent mosquito breeding habitats across other geographic regions, seasons, or ecological settings. A longitudinal study of the microbial communities of breeding habitats, spanning different seasons, could provide additional insights into the temporal dynamics of the microbiota in specific locations. The unequal sample sizes between natural and artificial habitats, in addition to the confounding of batch and biology, may have influenced differential abundance analyses. Moreover, only a subset of habitat-associated bacteria was functionally validated, while other ecological factors influencing oviposition, including physicochemical water properties, microbial interactions, predation, and interspecific competition, were not examined. Finally, the ecological mechanisms underlying habitat partitioning between *Aedes* and *Culex* mosquitoes remain to be elucidated.

## Conclusion

This study provides a comprehensive characterization of bacterial communities associated with the larval habitats of *Ae. aegypti* and demonstrates their potential role in shaping oviposition behavior. By integrating microbiome profiling, behavioral assays, and GC-MS analysis, we establish an association among habitat type (artificial vs natural), bacterial community composition, microbial metabolism, and mosquito oviposition behavior. These findings advance our understanding of mosquito-microbe chemical ecology and provide a foundation for developing bacterial volatile-based oviposition lures, surveillance tools, and environmentally sustainable vector control strategies. Future studies integrating longitudinal sampling, metabolomics, and field-based behavioral validation will further elucidate the temporal dynamics of habitat-associated microbial communities and their role in shaping mosquito population dynamics and disease transmission.

## Acknowledgements

TDD is the recipient of the Department of Science and Technology-Inspire Faculty Fellowship (Grant No. IFA-22-LSBM-264). MR is currently a PhD student at the Department of Biomedical Molecular Biology, Ghent University, Ghent, Belgium, the Causal Systems Immunology lab at the VIB-UGent Center for Inflammation Research, Ghent, Belgium and the VIB Center for AI & Computational Biology, Ghent, Belgium. We are thankful to ICMR-National Institute of Virology for infrastructural support and the Nanopore NGS facility for sequencing. We are also thankful to ICAR-National Research Centre for Grapes for the Sophisticated Analytical Instruments Facility (SAIF) for GC-MS support.

## Authors’ contributions

Conceived and designed the experiments: TDD, SPR, KSL, RV; Sample Collection: TDD, SPR, SS, SD, SD, SK; Performed the experiments: TDD, SS, SD, PK, AR; Analysed the data: TDD, MR, RV, SPR; Contributed reagents, materials, analysis tools: TDD, KSL, SPR, KB, RV, AY, KK; Wrote the paper: TDD, SPR, MR. Revised the manuscript: All authors read, revised, and approved the final version of the manuscript.

## Data availability statement

Metagenomic data for the water samples are deposited in https://www.ncbi.nlm.nih.gov/sra/PRJNA1513668

