## Supplementary Data for "Association between breeding-habitat bacterial communities and oviposition behavior of *Aedes aegypti:* Insights from arboviral outbreak-affected areas of Pune, Western India"

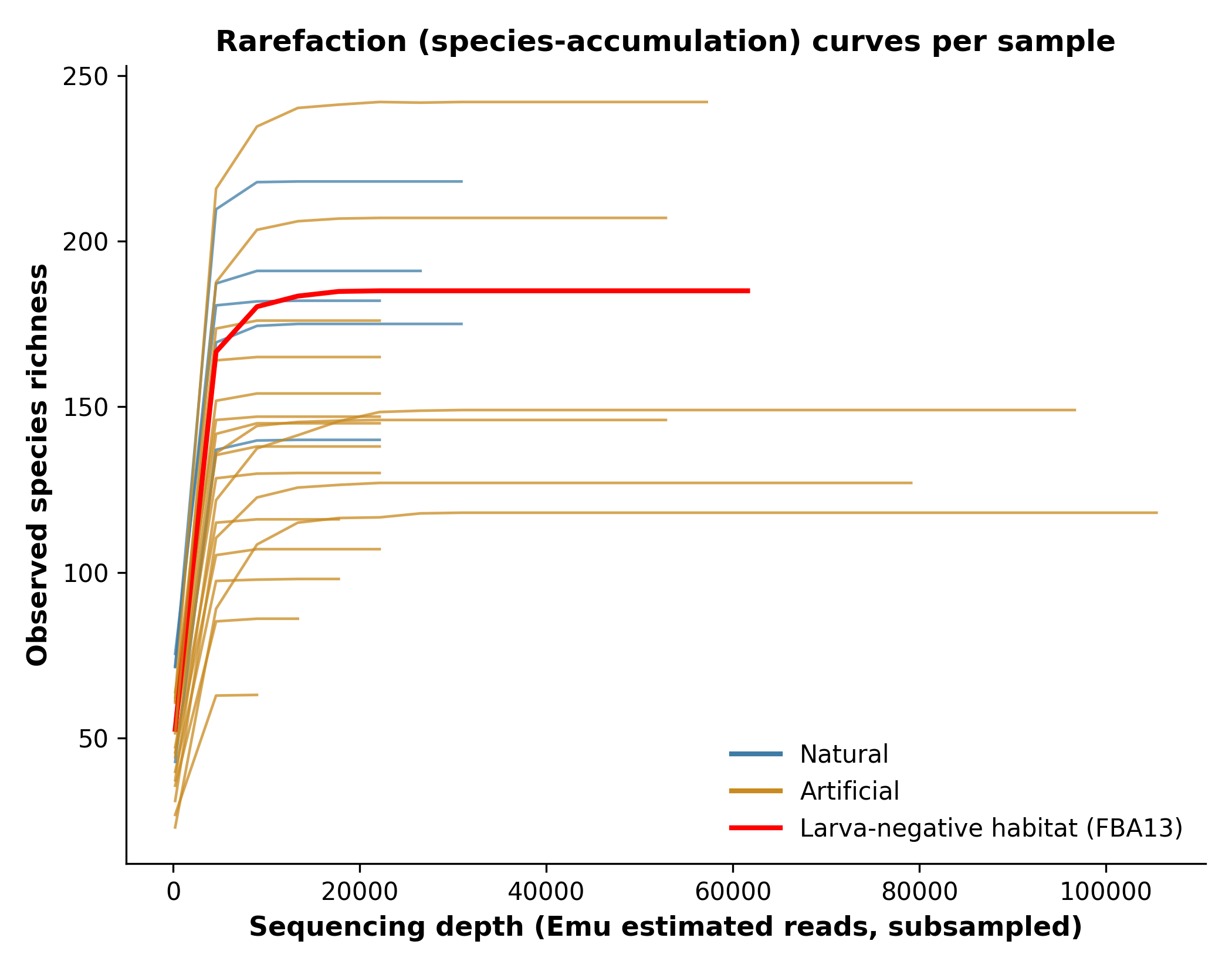

**Figure S1. Rarefaction sensitivity of the alpha-diversity result.** Per-sample rarefaction (species-accumulation) curves, colored by habitat type; the larva-negative artificial habitat (FBA13) is shown in red. All curves plateau well below the minimum observed depth (13,220 reads), indicating sequencing depth does not limit species detection at any sample's actual depth. Rarefying every biological sample to this minimum depth and recomputing Shannon/Simpson alpha diversity gave materially the same Wilcoxon rank-sum test result as the unrarefied analysis reported in the main text (Shannon: unrarefied P=0.019, rarefied P=0.0303; Simpson: unrarefied P=0.0028, rarefied P=0.0193; 100 rarefaction iterations per sample).

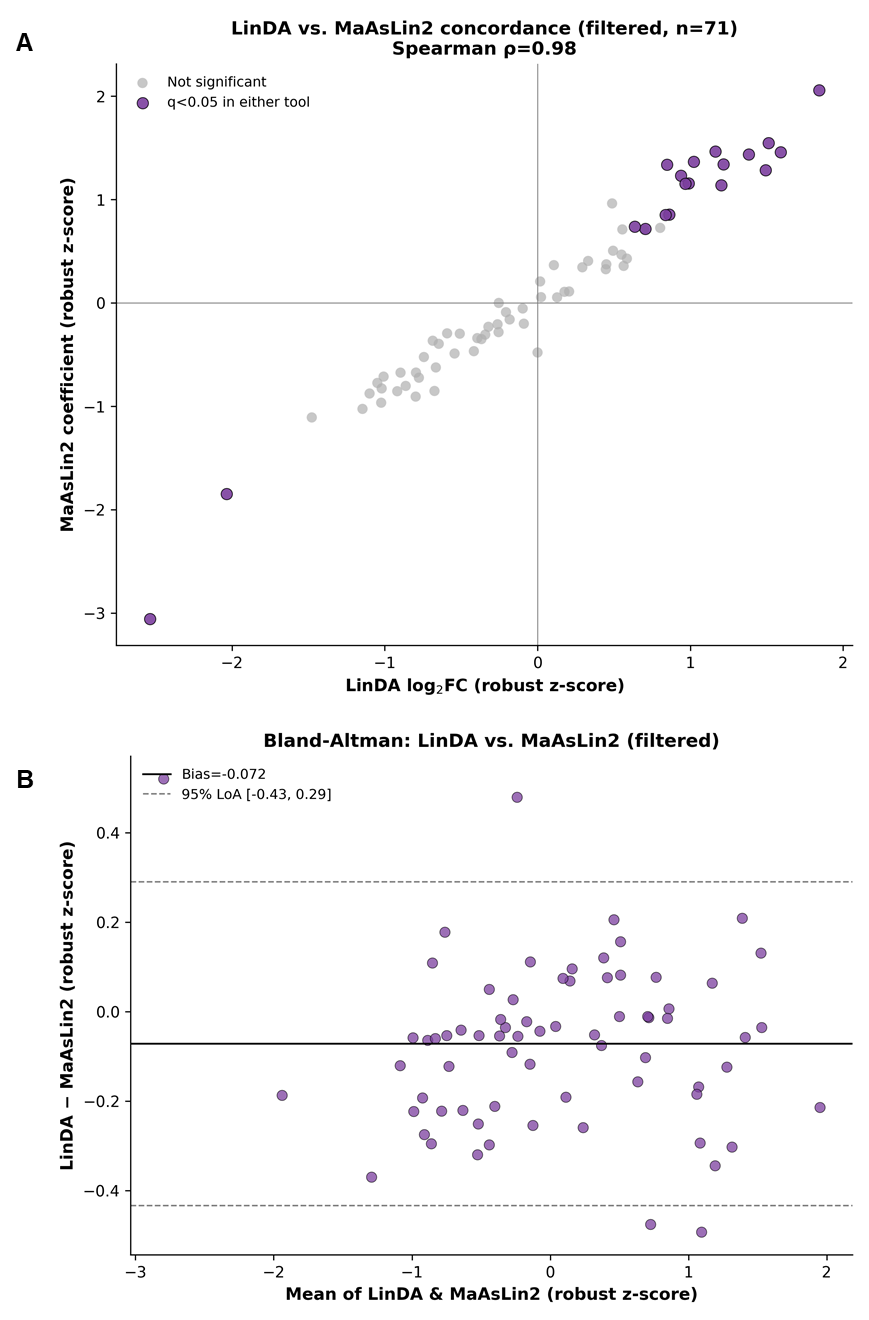

**Figure S2: LinDA vs. MaAsLin2 concordance (filtered species set)**. Cross-tool agreement between LinDA and MaAsLin2 on the 71 species passing the stringent filter (flow-cell-adjusted model, n=23 biological samples). Purple points: species significant (q<0.05) in either tool.

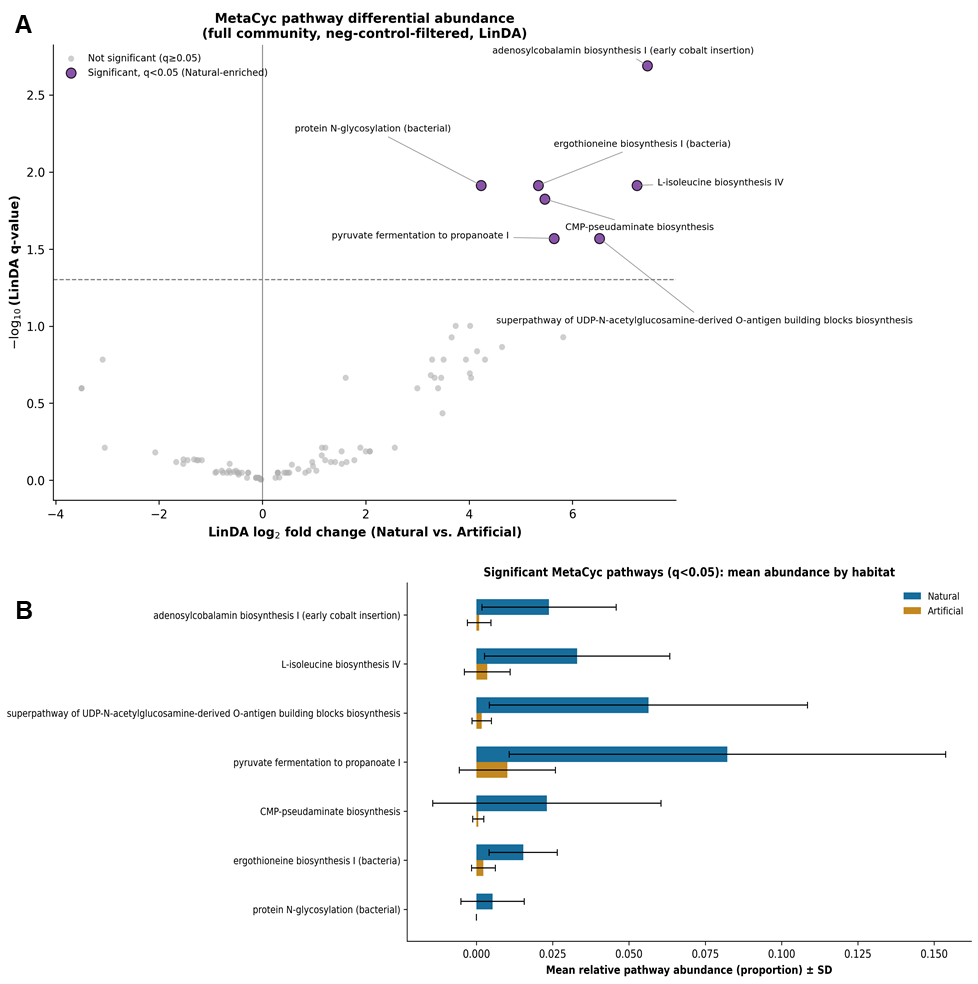

**Figure S3. MetaCyc pathway differential abundance full community.**

**Table S1: Details of sample collection site, type with sample codes used**

| **SL. No.** | **Sample Location** | **Sample Details** | **Sample code used for analysis**  **(Flow cell initials + barcode)** |
| --- | --- | --- | --- |
| 1. | Bibewadi | Artificial/plastic flower pot | FBA06 |
| 2. | Tathawade | Artificial/plastic bucket (Negative control) | FBA13 |
| 3. | Kharadi | Artificial/Basement | FBA14 |
| 4. | Shendurjane | Artificial/Flower pot | FBA18 |
| 5. | Pimpri | Artificial/Plastic drum | FBA19 |
| 6. | Vasavewadi | Artificial/Plastic drum | FBA22 |
| 7. | Pimpri | Artificial/School drum | FBA23 |
| 8. | Pashan | Natural/Lake | FAW02 |
| 9. | Alandi | Natural/River | FAW03 |
| 10. | Khadki | Artificial/Drum | FAW05 |
| 11. | Kala Kharak | Artificial/Drum | FAW06 |
| 12. | Pashan gaon | Artificial/Fridge | FAW07 |
| 13. | Katraj | Artificial/Tier | FAW08 |
| 14. | Susgaon | Artificial/Cattle Shed | FAW09 |
| 15. | Kopare | Artificial/Drum | FAW10 |
| 16. | Sutarwadi | Artificial/Drum | FAW11 |
| 17. | Aundh | Artificial/Flowering pot | FAW12 |
| 18. | Yerwada | Artificial Metal drum | FAW13 |
| 19. | MG road | Artificial/Black drum | FAW14 |
| 20. | Bopodi | Artificial/Bucket | FAW15 |
| 21. | Khadakwasla | Natural/Rock pool | FAW22 |
| 22. | Susgaon | Artificial/Plastic drum | FAW24 |
| 23. | Khadki | Natural/River | FAW04 |
| 24. | Kharadi Gaon | Natural/River | FAW01 |

**Table S2. Per-sample sequencing depth**

Sequencing depth is reported as the Emu estimated-count total (sum of per-species estimated counts) per sample, since Emu's own output does not include a separate raw-FASTQ read-count field; this is the same count basis used throughout the manuscript's differential-abundance analyses. Samples are sorted by depth.

| **Sample** | **Habitat type** | **Flow cell** | **Larva-negative habitat (FBA13)** | **Emu estimated-count total (depth)** |
| --- | --- | --- | --- | --- |
| FAW14 | Artificial | FAW | False | 13220 |
| FAW10 | Artificial | FAW | False | 17069 |
| FAW09 | Artificial | FAW | False | 18852 |
| FAW15 | Artificial | FAW | False | 22109 |
| FAW12 | Artificial | FAW | False | 22945 |
| FAW11 | Artificial | FAW | False | 22997 |
| FAW05 | Artificial | FAW | False | 23052 |
| FAW04 | Natural | FAW | False | 23323 |
| FAW06 | Artificial | FAW | False | 23482 |
| FAW08 | Artificial | FAW | False | 23506 |
| FAW24 | Artificial | FAW | False | 23740 |
| FAW07 | Artificial | FAW | False | 25818 |
| FAW13 | Artificial | FAW | False | 25897 |
| FAW22 | Natural | FAW | False | 26228 |
| FAW01 | Natural | FAW | False | 26873 |
| FAW03 | Natural | FAW | False | 31588 |
| FAW02 | Natural | FAW | False | 34677 |
| FBA22 | Artificial | FBA | False | 54251 |
| FBA19 | Artificial | FBA | False | 56979 |
| FBA06 | Artificial | FBA | False | 61031 |
| FBA13 | Artificial | FBA | True | 64760 |
| FBA14 | Artificial | FBA | False | 80724 |
| FBA23 | Artificial | FBA | False | 98361 |
| FBA18 | Artificial | FBA | False | 105420 |

**Table S3. Rarefied vs. unrarefied Shannon/Simpson diversity per biological sample.**

| **Sample** | **Habitat type** | **Shannon (unrarefied)** | **Shannon (rarefied)** | **Simpson (unrarefied)** | **Simpson (rarefied)** |
| --- | --- | --- | --- | --- | --- |
| FAW14 | Artificial | 1.993 | 1.992 | 0.648 | 0.648 |
| FAW10 | Artificial | 2.581 | 2.58 | 0.82 | 0.819 |
| FAW09 | Artificial | 3.074 | 3.073 | 0.903 | 0.903 |
| FAW15 | Artificial | 2.541 | 2.539 | 0.75 | 0.75 |
| FAW12 | Artificial | 3.935 | 3.932 | 0.96 | 0.96 |
| FAW11 | Artificial | 3.92 | 3.917 | 0.957 | 0.957 |
| FAW05 | Artificial | 3.439 | 3.438 | 0.927 | 0.927 |
| FAW04 | Natural | 4.313 | 4.309 | 0.976 | 0.976 |
| FAW06 | Artificial | 3.52 | 3.516 | 0.931 | 0.931 |
| FAW08 | Artificial | 3.604 | 3.602 | 0.948 | 0.948 |
| FAW24 | Artificial | 2.969 | 2.967 | 0.867 | 0.867 |
| FAW07 | Artificial | 3.551 | 3.548 | 0.934 | 0.934 |
| FAW13 | Artificial | 3.9 | 3.896 | 0.952 | 0.952 |
| FAW22 | Natural | 2.983 | 2.981 | 0.883 | 0.883 |
| FAW01 | Natural | 4.11 | 4.105 | 0.961 | 0.961 |
| FAW03 | Natural | 4.161 | 4.157 | 0.966 | 0.966 |
| FAW02 | Natural | 3.791 | 3.788 | 0.957 | 0.957 |
| FBA22 | Artificial | 3.399 | 3.396 | 0.94 | 0.94 |
| FBA19 | Artificial | 3.396 | 3.392 | 0.926 | 0.926 |
| FBA06 | Artificial | 3.893 | 3.885 | 0.955 | 0.955 |
| FBA14 | Artificial | 2.435 | 2.431 | 0.798 | 0.798 |
| FBA23 | Artificial | 2.988 | 2.984 | 0.891 | 0.891 |
| FBA18 | Artificial | 1.905 | 1.904 | 0.648 | 0.648 |

**Table S4. Primary candidate habitat-associated taxa: selection criteria (13 species)**

A species was designated a primary candidate habitat-associated taxon only if significant (q<0.05, Benjamini-Hochberg) in both LinDA and MaAsLin2, under the stringent combined filter (prevalence ≥20% of samples, mean relative abundance ≥0.1%, absent from the larva-negative artificial habitat FBA13), and in both condition A (flow-cell-adjusted, habitat_type + flow_cell, n=23) and condition B (single flow cell, FAW-only, habitat_type only, n=17).

| **Species** | **Direction** | **A: LinDA q** | **A: MaAsLin2 q** | **B: LinDA q** | **B: MaAsLin2 q** |
| --- | --- | --- | --- | --- | --- |
| *Arcobacter ellisii* | Natural | 1.06e-04 | 6.94e-04 | 1.10e-04 | 1.32e-02 |
| *Bacteroidales bacterium CF* | Natural | 1.06e-04 | 4.70e-04 | 2.70e-04 | 1.28e-02 |
| *Arcobacter cryaerophilus* | Natural | 2.03e-04 | 2.45e-03 | 6.85e-04 | 1.32e-02 |
| *Methyloparacoccus murrellii* | Natural | 2.03e-04 | 6.14e-03 | 1.83e-04 | 1.81e-02 |
| *Novosphingobium naphthae* | Natural | 1.11e-03 | 1.91e-03 | 1.13e-03 | 1.32e-02 |
| *Enterobacter asburiae* | Artificial | 1.67e-03 | 9.46e-05 | 5.67e-03 | 1.32e-02 |
| *Rhodobacter capsulatus* | Natural | 1.67e-03 | 1.43e-02 | 1.23e-03 | 3.54e-02 |
| *Enterobacter roggenkampii* | Artificial | 9.52e-03 | 2.45e-03 | 1.54e-02 | 1.32e-02 |
| *Bordetella sp. HZ20* | Natural | 9.74e-03 | 2.75e-02 | 1.54e-02 | 4.90e-02 |
| *Dechloromonas agitata* | Natural | 9.74e-03 | 1.57e-02 | 1.49e-02 | 3.54e-02 |
| *Hydrogenophaga bisanensis* | Natural | 9.74e-03 | 1.43e-02 | 8.93e-03 | 2.55e-02 |
| *Achromobacter panacis* | Natural | 1.74e-02 | 4.70e-02 | 7.32e-03 | 3.54e-02 |
| *Quisquiliibacterium transsilvanicum* | Natural | 2.34e-02 | 7.41e-03 | 2.11e-02 | 1.32e-02 |

**Table S5: Significant MetaCyc pathways (full community, neg-control-filtered, LinDA, q<0.05)**

| **MetaCyc ID** | **Pathway** | **LinDA log2FC (Nat. vs Art.)** | **q-value** | **Primary-candidate contributors** |
| --- | --- | --- | --- | --- |
| P108-PWY | pyruvate fermentation to propanoate I | 5.64 | 2.69e-02 | none of the 13 primary candidates |
| PWY-5104 | L-isoleucine biosynthesis IV | 7.25 | 1.22e-02 | none of the 13 primary candidates |
| PWY-5507 | adenosylcobalamin biosynthesis I (early cobalt insertion) | 7.45 | 2.04e-03 | none of the 13 primary candidates |
| PWY-6143 | CMP-pseudaminate biosynthesis | 5.46 | 1.50e-02 | Arcobacter cryaerophilus (72%) |
| PWY-7031 | protein N-glycosylation (bacterial) | 4.23 | 1.22e-02 | none of the 13 primary candidates |
| PWY-7255 | ergothioneine biosynthesis I (bacteria) | 5.34 | 1.22e-02 | none of the 13 primary candidates |
| PWY-7332 | superpathway of UDP-N-acetylglucosamine-derived O-antigen building blocks biosynthesis | 6.52 | 2.69e-02 | none of the 13 primary candidates |

“Primary-candidate contributors” lists which of the 13 batch-robust primary candidate species (Figure 5) contribute to each pathway, with the mean fraction of that pathway's predicted abundance attributable to that taxon in samples where it contributes at all (PICRUSt2 stratified output).
